# Performance reserves in brain-imaging-based phenotype prediction

**DOI:** 10.1101/2022.02.23.481601

**Authors:** Marc-Andre Schulz, Danilo Bzdok, Stefan Haufe, John-Dylan Haynes, Kerstin Ritter

## Abstract

Machine learning studies have shown that various phenotypes can be predicted from structural and functional brain images. However, in most such studies, prediction performance ranged from moderate to disappointing. It is unclear whether prediction performance will substantially improve with larger sample sizes or whether insufficient predictive information in brain images impedes further progress. Here, we systematically assess the effect of sample size on prediction performance using sample sizes far beyond what is possible in common neuroimaging studies. We project 3-9 fold improvements in prediction performance for behavioral and mental health phenotypes when moving from one thousand to one million samples. Moreover, we find that moving from single imaging modalities to multimodal input data can lead to further improvements in prediction performance, often on par with doubling the sample size. Our analyses reveal considerable performance reserves for neuroimaging-based phenotype prediction. Machine learning models may benefit much more from extremely large neuroimaging datasets than currently believed.

## Introduction

Advances in neuroimaging have provided unprecedented insight into the structure and function of the human brain. Concurrently, high-resolution brain scans have become increasingly cost-effective and have raised hopes for automated disease diagnoses and clinical endpoint prediction based on neuroimaging data. Significant progress has been made in some areas of application, for instance in the automated detection of brain atrophy (Jack et al., 2004; Plant et al., 2010; Rocca et al., 2017) and segmentation of brain lesions (Kamnitsas et al., 2017; Akkus et al., 2017). However, the prediction of cognitive and behavioral phenotypes and the diagnosis of psychiatric diseases has remained challenging (Kapur et al., 2012; Woo et al., 2017). Whether such challenges in neuroimaging-based phenotype prediction are primarily due to insufficient sample sizes or a lack of predictive information in neuroimaging data remains to be clarified.

Moving from group-level inference to accurate single subject prediction and searching for intricate patterns in high-dimensional data can require sample sizes that are orders of magnitude larger than those of traditional neuroimaging studies (Arbabshirani et al., 2017; Bzdok et al., 2019). Indeed, neuroimaging datasets have continuously grown in sample size over the last decade (Szucs & Ioannidis, 2020). While early neuroimaging studies were limited to only tens of participants, researchers now often include hundreds of participants aggregated from multiple acquisition sites. Large-scale data collection initiatives have grown from the Human Connectome Project (Van Essen et al., 2013), with roughly one thousand participants, to the UK Biobank’s Imaging Initiative (Littlejohns et al., 2020; Miller et al., 2016), with currently 46 thousand participants and an end-goal of one hundred thousand participants. Other researchers have proposed a Million Brains Initiative to facilitate precision medicine in the USA (Liebeskind et al., 2017). Since collecting large neuroimaging datasets is an elaborate and expensive endeavor, we ask: What sample size is really necessary for reliable single subject prediction of cognitive and behavioral phenotypes from brain images?

At the same time, there exists some controversy about how much predictive information can plausibly be extracted from conventional neuroimaging data. While high prediction accuracy has been reported for some phenotypes (overview in Arbabshirani et al., 2017), these findings have generated controversial debate (Flint et al., 2021; Varoquaux, 2018; Woo et al., 2017), and reliable neuroimaging biomarkers for psychiatric disease have remained largely elusive (Calhoun et al., 2021; Kapur et al., 2012; Woo et al., 2017). High researcher degrees of freedom combined with imperfect model validation practices may have inflated results in small-sample studies (Flint et al., 2021; Poldrack et al., 2020). In line with this hypothesis of over-optimistic reporting, meta-analyses have shown a pronounced inverse trend between prediction accuracy and sample size (Neuhaus & Popescu, 2018). Even under ideal circumstances, it is unclear to what extent intricate cognitive or behavioral traits can be predicted based on brain images (Uttal, 2011; Woo et al., 2017). Conventional neuroimaging could operate at impotent levels of abstraction or spatiotemporal resolution (Logothetis, 2008; Uttal, 2011). Moreover, neuroimaging is particularly affected by high levels of noise in the data (e.g., in functional magnetic resonance imaging [MRI] where the phenomenon being studied often only makes up a small part of the blood-oxygen-level-dependent signal; Liu, 2016; Raz et al., 2005) and in the target labels (e.g., inherent subjectivity of symptoms, low retest-reliability of diagnoses; Hyman, 2007; Kraemer et al., 2012; Regier et al., 2013). A lack of predictive information in the data or high levels of noise can pose an upper limit to the prediction accuracy, even in the limit of infinite samples and perfect machine learning algorithms. This raises the question: Do structural and functional brain images contain enough exploitable predictive information to be useful for precision medicine?

The two questions of sample size and exploitable predictive information are closely related. If there was insufficient predictive information in the data, then adding more participants would not improve prediction accuracy and there would be no need for a Million Brain Project. Conversely, if we find that increasing the sample size yields continuous improvements in predictive accuracy, then we can conclude that we have not yet exhausted the predictive information contained in the data and there is hope for accurate single subject prediction. Therefore, it is crucial to clarify whether low prediction accuracy in a small or medium-sized study primarily reflects fundamental limitations on predictive information (related to the so-called irreducible error, independent of sample size) or whether accuracy can be improved by increasing the sample size. This translates into asking: Can we mathematically characterize the empirical relationship between sample size and achievable prediction accuracy - the “learning curve” - for a given target phenotype? Characterizing and extrapolating such scaling laws would answer both original questions: It would provide estimates of the necessary sample size to reach a certain accuracy level as well as estimates of the highest achievable accuracy for a target phenotype given infinite samples.

Theoretical results from statistical learning theory state that the prediction accuracy typically scales as a power law function of the sample size (Amari, 1993; Amari et al., 1992; Haussler et al., 1996; Hutter, 2021). Empirically, power law scaling of learning curves has been shown for models ranging from linear estimators to deep neural networks (Cortes et al., 1994; Hestness et al., 2017). Hence, estimating power law parameters from an empirical learning curve allows one to extrapolate the learning curve beyond the available sample size and thereby to delineate two key properties of the given prediction task: First, the convergence point of the learning curve represents the maximally achievable prediction accuracy. This can be taken as an estimate (technically a lower bound, see Discussion) of the exploitable predictive information encoded in the data. Second, the speed of convergence can be defined as the sample efficiency, i.e., the amount of data needed by the model to learn the task. Intuitively, the former represents how easy it is to *make an accurate prediction* while the latter reflects how easy it is to *learn how to predict*. For a given brain imaging modality and phenotype, the power law scaling behavior of the learning curve should allow one to infer both the maximally achievable prediction accuracy and the necessary sample size to achieve clinically useful performance, thus quantitatively engaging two core aspects of feasibility for precision medicine.

In sum, deriving realistic estimates of achievable prediction accuracy and required sample sizes is crucial to gauge whether a predictive modeling approach may be suitable for single subject prediction in precision medicine. Here, we systematically evaluate learning curves for different neuroimaging data modalities (structural and functional MRI) and a diverse set of demographic, cognitive, behavioral, and mental health phenotypes. We assess the validity of extrapolated learning curves and analyze which representations of brain imaging data have the highest predictive potential in the limit of large sample sizes and thus delineate areas of feasibility and infeasibility in the landscape of machine learning empowered predictions in precision medicine.

## Results

We based our analyses on the UK Biobank brain-imaging dataset, which is the largest uniformly acquired brain-imaging dataset available to date (46197 participants, June 2021 release; Miller et al., 2016). Given its scope and extensive quality control, we argue that the UK Biobank can be considered a current “best-case scenario” for the upper end of available neuroimaging data analysis. The UK Biobank provides imaging-derived-phenotypes (IDPs) of T1-weighted structural brain MRI (regional gray and white matter volumes, cortical thickness and surface area), resting-state functional brain MRI (rfMRI; ICA-based functional connectivity), and diffusion-weighted imaging (DWI; anisotropy and diffusivity measures). From these IDPs, we predicted widely studied phenotypes from sociodemographic (age, sex, education score, household size), cognitive (fluid intelligence, reaction time, numeric memory, trail making), behavioral (alcohol, tobacco, and TV consumption, physical activity), and mental health (financial and friendship satisfaction, depression, neuroticism) domains. We used regularized linear models to predict target phenotypes from brain imaging data. Such models are considered highly competitive in the analysis of neuroimaging data and performed on par with more complex nonlinear models on similar prediction tasks (Bzdok & Ioannidis, 2019; Dadi et al., 2019; Dufumier et al., 2021; Schulz et al., 2020). To derive rigorous learning curves for our prediction scenarios, we repeatedly pulled smaller subsamples from the UK Biobank data (n=256, 362, 512, …, 32k) and trained and evaluated our models for each combination of sample size, input modality, and target phenotype. For each resulting learning curve, we fitted a standard power law [α n^-β^ + γ] to the empirical measures (cf. Cortes et al., 1994; Hutter, 2021). For details on brain imaging data, target phenotypes, machine learning models, and evaluation procedure, please refer to the Methods section.

### Learning curves follow power laws

To assess how accurately the power law function family describes learning curves for neuroimaging-based phenotype prediction, we calculated goodness-of-fit statistics (R^2^, χ^2^) for each of the 48 (16 target phenotypes x 3 modalities) prediction tasks. An average coefficient of determination R^2^ of 0.990 (SD=0.015, min=0.902) indicated an excellent fit between power law and empirical learning curves (Fig. 1). Reduced χ^2^ statistics, on average 0.035 (SD=0.038, max=0.207), were fully compatible with our power law hypothesis. Note that χ^2^<<1, suggesting that our estimated measurement uncertainties were rather conservative (Bevington & Robinson, 2002). Average goodness-of-fit statistics were comparable between imaging modalities (T1 R^2^=0.994, rfMRI R^2^=0.986, DWI R^2^=0.991) and target-phenotype categories (sociodemographic R^2^=0.993, cognitive R^2^=0.996, behavioral R^2^=0.982, mental health R^2^=0.992). The observed scaling behavior of prediction accuracy with increasing sample size closely followed a power law for all investigated target phenotypes and imaging modalities.

**Figure 1:**
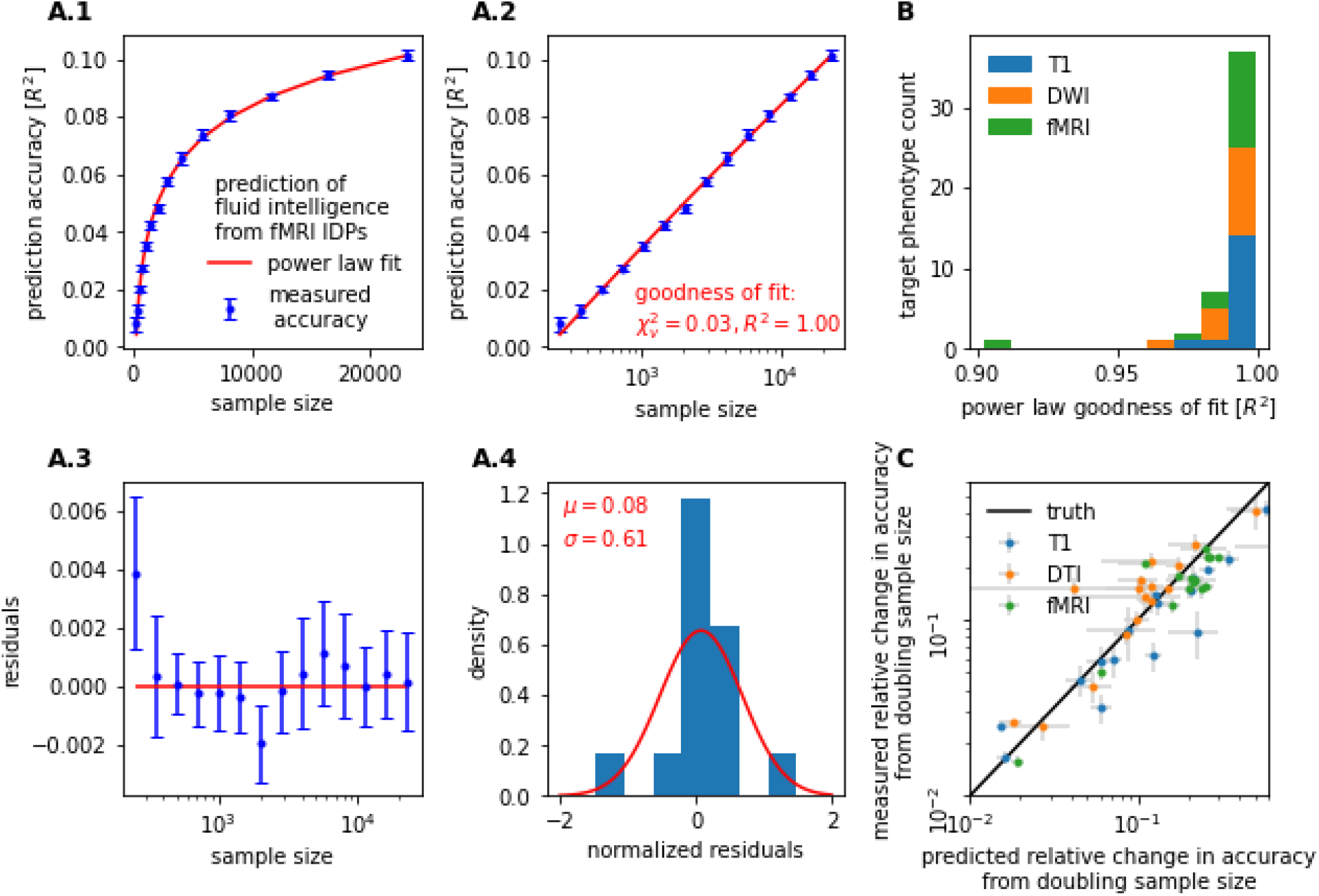
Learning curves for neuroimaging-based phenotype prediction precisely follow a power law function. Prediction accuracy scales with the number of training samples. The precise nature of this relationship can be described by a simple power law [α n^-β^ + γ]. A.1) For instance, when predicting fluid intelligence from rfMRI data using ridge regression, out-of-sample accuracy (blue) closely followed the fitted power law (red). A.2) We observed stable and continuous improvements in accuracy with increasing sample size, i.e., approximately linear scaling of prediction accuracy with log(n). A.3-4) Residuals of the power law fit gave no indication of systematic deviations between measured accuracy and fitted power law. B) Power law scaling was observed in all evaluated prediction tasks (i.e., combinations of imaging modality and target phenotype), with a goodness-of-fit R^2^ between measured learning curve and power law of on average 0.990 (SD=0.015, min=0.902) C) Learning curve extrapolation predicted accuracy achievable on unseen larger samples. Shown are projected gains in prediction accuracy derived from learning curve extrapolation on the y-axis in relation to observed gains in prediction accuracy on the x-axis. Both were derived by doubling the training sample size from eight thousand to 16 thousand. Error bars indicate standard error of the mean (SEM).

Once a power law had been fitted to a learning curve, the curve could be mathematically extrapolated beyond the available sample size. To validate the results of such extrapolations, we fitted our power laws exclusively on sample sizes of 256 to eight thousand, retaining the doubled sample size of 16 thousand as a test set. We evaluated the out-of-sample extrapolation by comparing the extrapolated gain in prediction accuracy from eight thousand to 16 thousand samples to the ground-truth gain (Fig. 1-C). Extrapolation and ground-truth pairs were aggregated for each of our prediction tasks. Power law extrapolations of the learning curve were highly accurate with an average coefficient of determination of R^2^=0.788 (T1 R^2^=0.828, rfMRI R^2^=0.716, DWI R^2^=0.710). The obtained goodness-of-fit statistics and out-of-sample extrapolation suggest that learning curve power laws can be used to estimate maximally achievable accuracy as well as necessary sample sizes for threshold accuracies.

### Learning curve extrapolation reveals performance reserves in phenotype prediction

All examined target phenotypes could be predicted from our T1, rfMRI, and DWI brain data, although to varying degrees of accuracy. Prediction accuracy (classification accuracy or regression R^2^ respectively) was highest for sex and age (best performing modality: accuracy / R^2^ at maximum sample size; sex - T1: 0.968±0.001, age - T1: 0.762±0.002). Sociodemographic phenotypes (education score - DWI : 0.034±0.002, household size - T1 : 0.085±0.004) and cognitive phenotypes (reaction time - T1 : 0.090±0.002, numeric memory - rfMRI : 0.070±0.002, fluid intelligence - rfMRI : 0.101±0.002, trail making - T1 : 0.120±0.003) scored higher than behavioral phenotypes (alcohol - rfMRI : 0.063±0.003, smoking T1 : 0.048±0.002, TV consumption - rfMRI : 0.077±0.002, physical activity - T1 : 0.011±0.001) and mental health phenotypes (depression - rfMRI : 0.564±0.001, neuroticism - T1 : 0.030±0.001, financial satisfaction - T1 : 0.023±0.001, friendship satisfaction - rfMRI : 0.023±0.002). However, different target phenotypes yielded wildly heterogeneous learning curves (Fig. 2). For sex and age, all three neuroimaging modalities showed saturation of prediction accuracy when increasing the sample size beyond 16 thousand. In contrast, for every other target phenotype, at least one modality showed stable and continuous improvements in accuracy with increasing sample size (i.e., approximately linear scaling of prediction accuracy with log(n), see Fig. 1-A.2) up to the 32 thousand training samples from the UK Biobank. In these data analysis settings, learning curve extrapolation projected continuous improvements up to at least one million samples.

**Figure 2:**
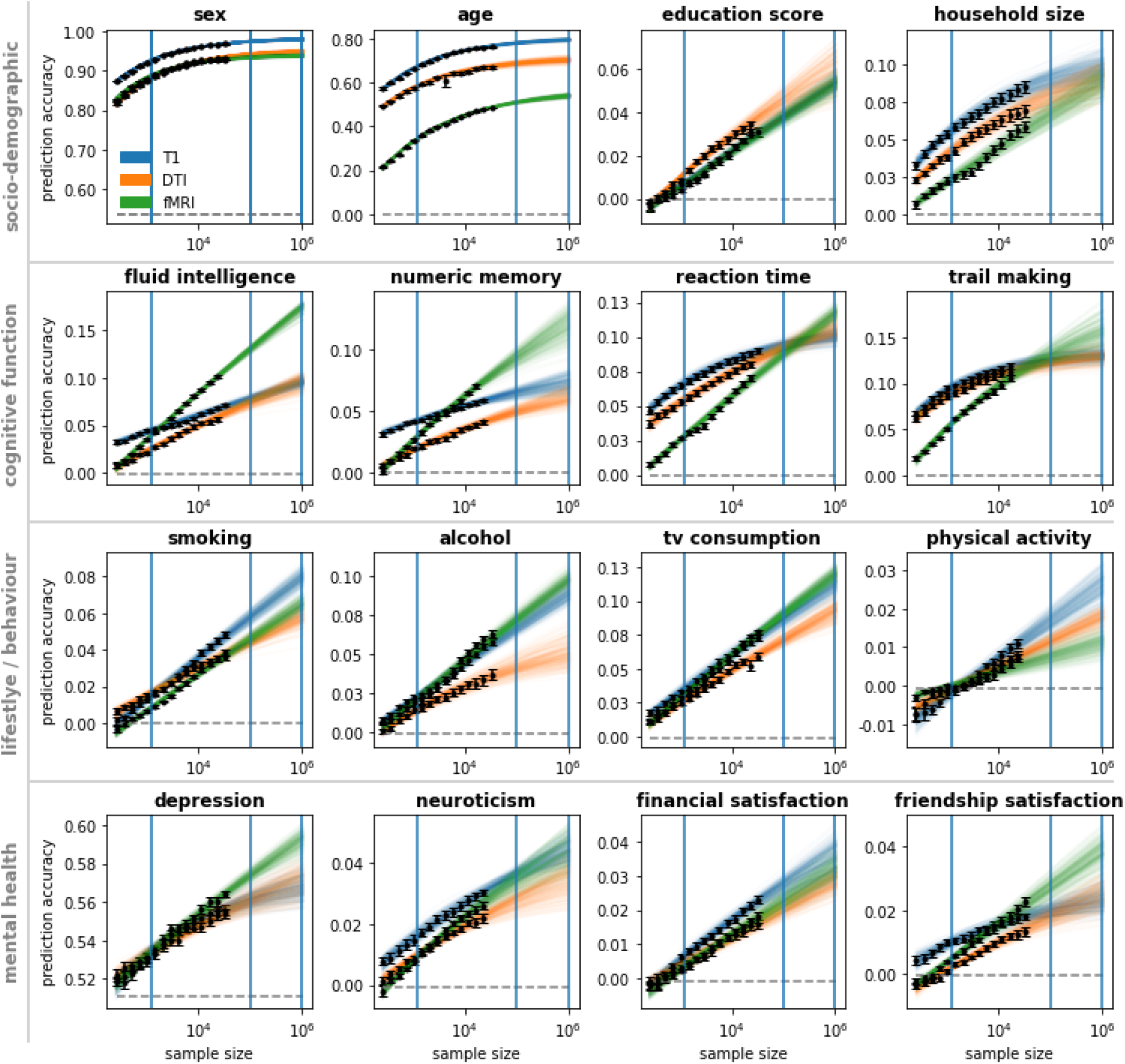
Linear models are operating far below ceiling accuracy for most target phenotype predictions. Learning curves show the collective results obtained from regularized linear models using T1, DWI, and rfMRI data to predict sociodemographic, cognitive function, behavior/lifestyle, and mental health phenotypes. Training datasets were subsampled from the UK Biobank up to a size of 32 thousand participants. Learning curves were extrapolated beyond 32 thousand participants. To indicate extrapolation uncertainty, each colored line represents a power law fit based on a bootstrap sample of observed accuracies. Observed prediction accuracies are marked black; majority classifier / median regression baselines are marked dashed gray. Blue vertical lines indicate the sample size of the Human Connectome Project (1k), the imaging sample size goal of the UK Biobank (100k), and the proposed Million Brain Initiative (1M). Error bars indicate SEM.

To quantify how prediction performance is projected to gain from further increases in sample size beyond our sample, we used the Human Connectome Project sample size (1k) as a reference and calculated the expected relative change in accuracy when escalating from one thousand (Human Connectome Project) to one million (Million Brain Project goal) samples (Fig. 3). The largest relative change was projected for mental health phenotypes (friendship satisfaction - DWI : 8.89±2.14, financial satisfaction - rfMRI : 8.34±1.47, depression - rfMRI : 2.15±0.27, neuroticism - rfMRI : 3.99±0.58) followed by behavioral phenotypes (alcohol – rfMRI : 3.94±0.23, smoking rfMRI : 7.74±0.97, TV consumption - rfMRI : 3.13±0.18, physical activity could not be reliably estimated due to near-zero baseline), cognitive phenotypes (reaction time - rfMRI : 3.35±0.18, numeric memory - rfMRI : 3.41±0.47, fluid intelligence - rfMRI : 3.92±0.19, trail making - rfMRI : 1.92±0.21), and sociodemographic phenotypes (household size - rfMRI : 3.04±0.27, education - T1 : 6.61±0.71). Consistent with observed learning curve saturation, sex (DWI : 0.21±0.01) and age (rfMRI : 0.62±0.03) age scored lowest. In other words, we project a 3-9 fold increase in prediction performance for behavioral and mental health phenotypes when moving from one thousand to one million samples. Our results suggest that for most investigated target phenotypes, linear models still operate far below their respective performance ceilings at currently available sample sizes.

**Figure 3:**
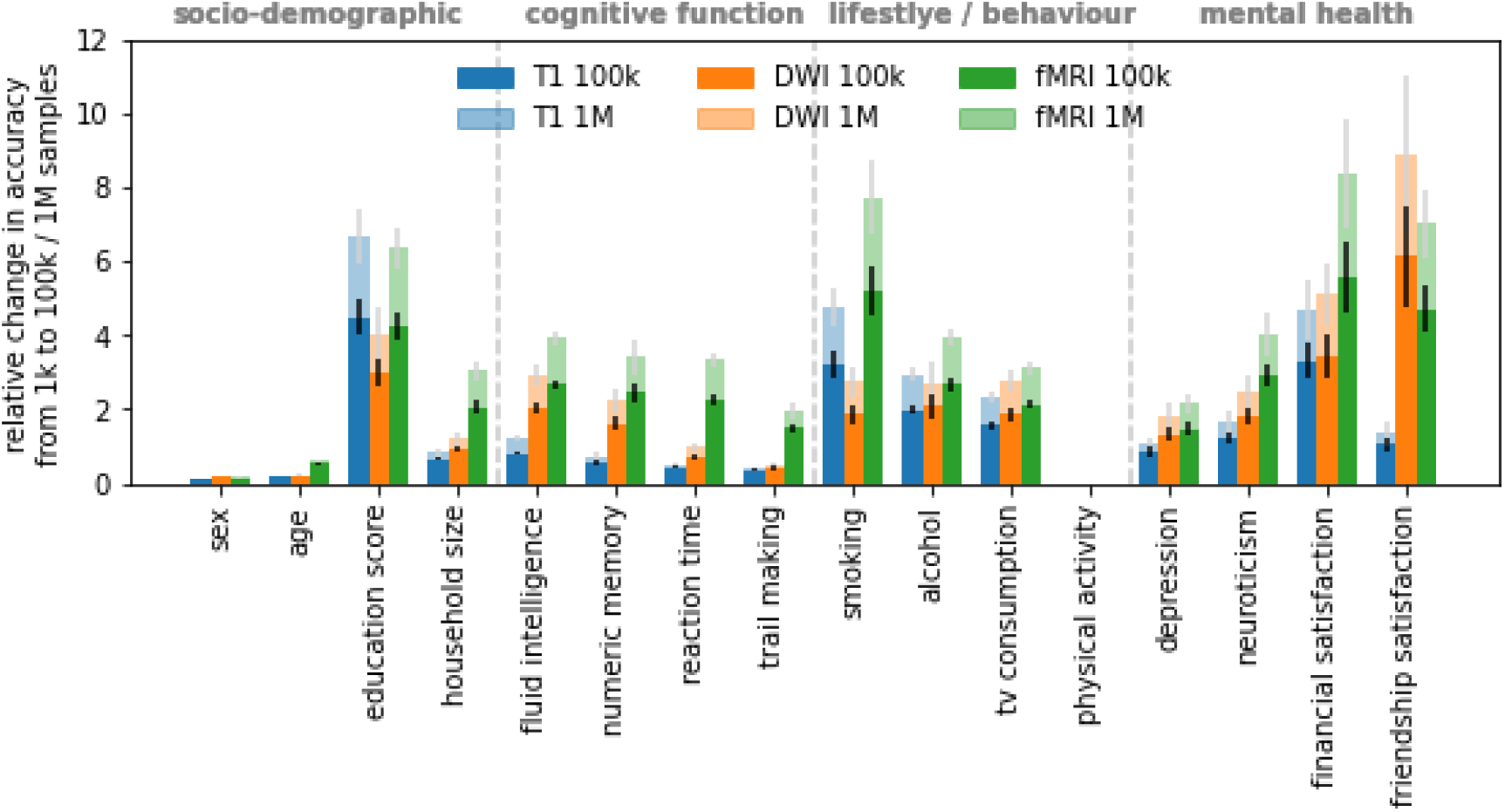
Multi fold-gains in prediction performance are projected for behavioral and mental health phenotypes when moving from one thousand to one million samples. Shown is the relative increase in prediction accuracy per modality and target phenotype derived from learning curve extrapolation on regularized linear models. Results for physical activity could not be reliably estimated due to near-zero baseline (cf. Fig. 2). Error bars indicate SEM.

### Optimal choice of imaging modality depends on both target phenotype and sample size

The scaling trajectory of prediction performance with sample size expressed a multitude of patterns based on the specific combination of evaluated brain-imaging data and target phenotypes (Fig. 2). No single modality consistently outperformed the other modalities, nor did we observe a consistent rank order of modalities over the set of prediction targets. Even for a single target phenotype, different modalities expressed different scaling behavior so that the accuracy hierarchy of modalities would often change with increasing sample size. Nearly every possible rank order of modalities was observed in at least one target phenotype and sample size range. The most frequent rank order was T1 > DWI > rfMRI, representing 40.26 % of target x sample size combinations excluding extrapolated values. However, extrapolated to one million samples, rfMRI was projected to outperform T1 and DWI for the majority (9/16) of target phenotypes. A compelling example is the case of fluid intelligence (Fig. 2), where T1 outperformed rfMRI by 2.47 percentage points for small sample sizes (n=256) but was projected to be outperformed by rfMRI by 7.92 percentage points for very large sample sizes (n=1M). In some but not all of such cases, T1 and DWI approached saturation accuracy (cf. cognitive function, Fig. 2). In contrast, for rfMRI, learning curve extrapolation predicted continuous improvements up to at least one million samples (except for sex and age). In sum, the best performing modality depended on both target phenotype and sample size, leading to pronounced modality cross-over effects for some target phenotypes. Further, extrapolation of performance scaling with increasing sample size suggests a higher accuracy ceiling for rfMRI than for T1 and DWI.

### Multimodal data substantially boosts prediction performance

Does combining different imaging modalities yield improvements in out-of-sample prediction performance over a single-modality baseline? Information extracted from different imaging modalities might be independent, so that combining modalities would improve out-of-sample prediction performance, or redundant, so that combining modalities would be ineffective. To investigate the impact of multimodal data, we concatenated the imaging data into dual-modality and triple-modality feature spaces, and retained the first 512 principal components for each respective feature space to align feature dimensionalities (cf. Abrol et al., 2021; Schulz et al., 2020). Phenotype prediction based on the combination of T1, DWI, and rfMRI data outperformed prediction based on the respective best single modality for all target phenotypes and led to an average relative increase in accuracy of 30.78% (SD=18.57 p.p.) at 16 thousand samples. Switching from single modalities to multimodal input data led to improvements in prediction accuracy on par with doubling the sample size from eight thousand to 16 thousand, for 10 out of 16 target phenotypes. Learning curves observed in single-modality experiments (Fig. 2) were mirrored in our analysis of multimodal feature spaces (Fig 4). Our results suggest that different brain imaging modalities do not simply reflect the same limited set of variables but instead offer complementary non-redundant predictive information for the majority of target phenotypes.

**Figure 4:**
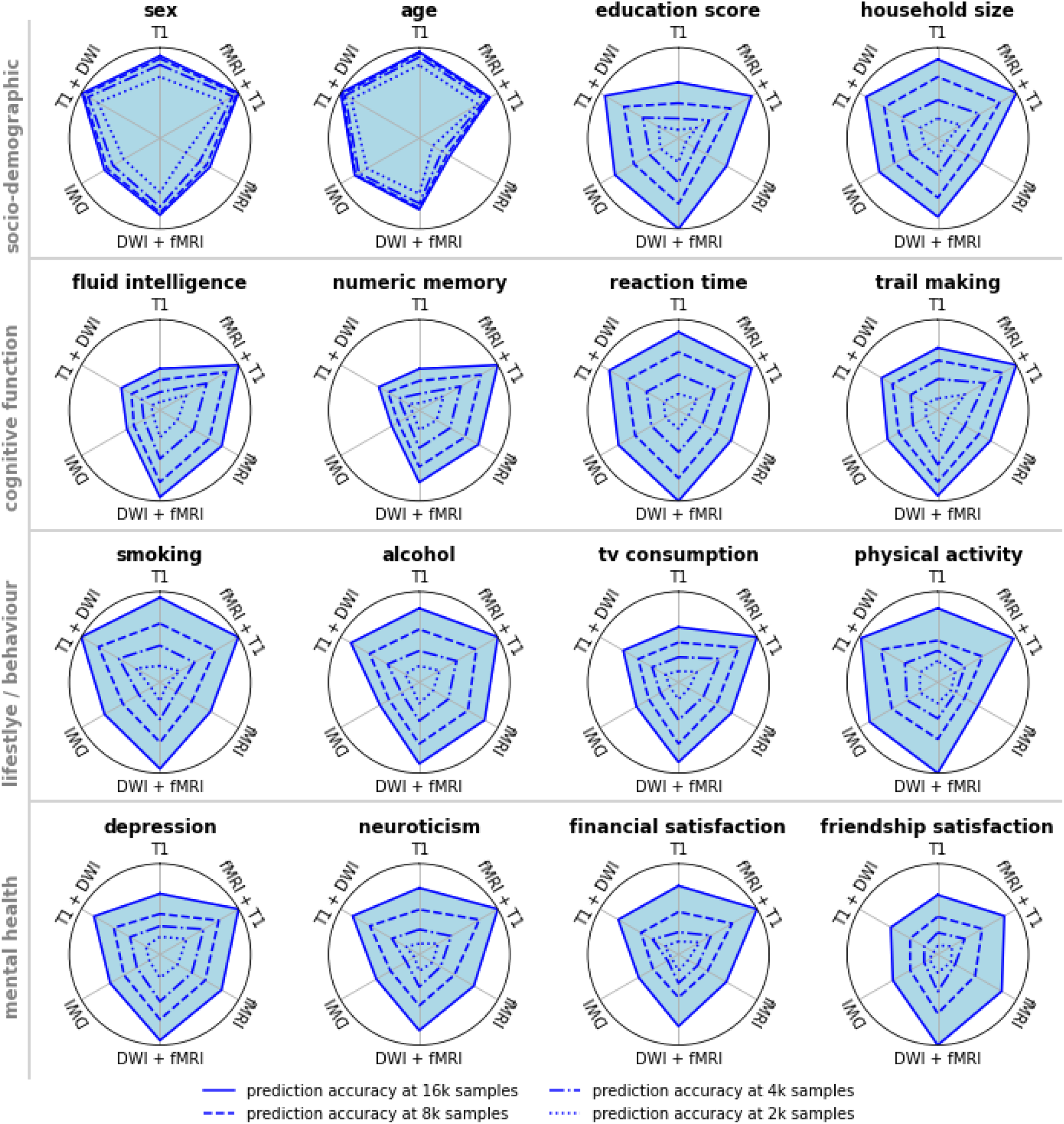
Augmenting single-modality feature spaces to incorporate multimodal input data can lead to improvements in prediction accuracy on par with doubling the sample size. The 512 leading principal components of single-modality data, or of concatenated dual-modality data, were used as the basis for phenotype prediction. Pictured is the min-max scaled prediction accuracy, with accuracy at one thousand training samples representing the origin of the respective graph. Switching from single modalities to multimodal input data led to improvements in prediction accuracy for all target phenotypes. For 10 out of 16 target phenotypes, improvements from multimodality were comparable to improvements from doubling the sample size from eight thousand to 16 thousand. Different brain imaging modalities appear to provide complementary, non-redundant predictive information for most target phenotypes.

### Direct comparison of linear and nonlinear models

In recent work (Schulz et al., 2020), we observed that linear and more expressive nonlinear machine learning models did not show relevant differences in performance for sex and age prediction based on T1 and rfMRI data for up to eight thousand training samples. We replicated these analyses for age, sex, fluid intelligence, and depression and on up to 32 thousand training samples, comparing linear ridge regression with its nonlinear counterpart RBF-kernelized ridge regression. Only in sex and age prediction based on DWI with large sample sizes above 16 thousand, nonlinear models appeared to marginally outperform their linear counterparts (Fig. 5-A.1). On all other evaluated prediction settings, linear models performed on par with nonlinear machine learning models. We found no consistent evidence of exploitable predictive nonlinear structure in neuroimaging data. Our collective results did not qualitatively differ between linear and nonlinear machine learning models.

**Figure 5:**
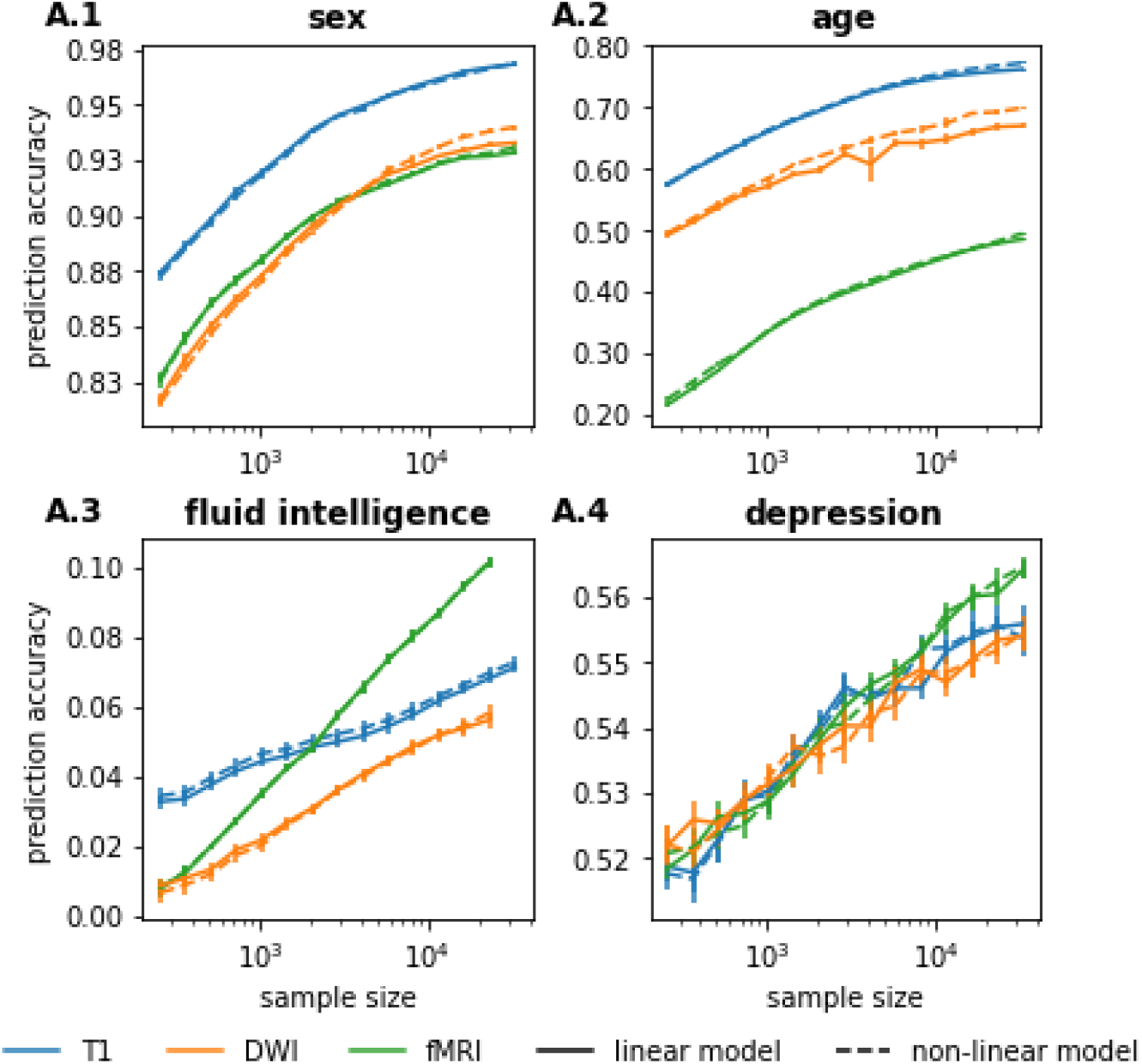
Linear models performed on par with nonlinear machine learning models in neuroimaging-based phenotype prediction. We found no consistent evidence of exploitable predictive nonlinear structure in neuroimaging data. Only for DWI-based prediction of sex and age at large (>16k) training sample sizes did nonlinear models marginally outperform their linear counterparts. Pictured are results for linear and RBF-kernelized nonlinear ridge regression. For other nonlinear machine learning models, see Supplementary Material. Error bars indicate SEM.

## Discussion

Current research on neuroimaging-based precision medicine suffers from an information vacuum. Without principled estimates on the mutual information between brain imaging data and a given target phenotype, researchers are limited to a trial and error approach in which they are left guessing whether a machine learning model’s bad performance is due to technical error, insufficient sample size, or lack of predictive information in the data. Principled estimates on the mutual information between brain imaging data and target phenotype should help streamline the development of complex machine learning models, inform large-scale data collection initiatives, and help allocate resources to the most promising phenotypes.

In the present study, we introduced learning curve extrapolation as an effective tool to estimate achievable prediction accuracy and sample size requirements for neuroimaging-based phenotype prediction. Our systematic characterization of learning curves for different imaging modalities and target phenotypes revealed three major findings: First, for most of the investigated target phenotypes, prediction performance continued to improve with additional samples, even at the limit of currently available sample sizes. These untapped performance reserves suggest that machine learning models in neuroimaging-based phenotype prediction operate far below their ceiling accuracy, raising hopes for accurate single subject prediction in precision medicine. Second, different imaging modalities yielded unique predictive information, and combining modalities led to improvements in prediction accuracy on par with doubling the sample size. Moving from single imaging modalities to multimodal input data may unlock further substantial performance reserves for neuroimaging-based phenotype prediction. Finally, a majority of target phenotypes exhibited cross-over effects with regard to the best performing modality. Instead of one single imaging modality being optimal for predicting a given target phenotype, the best performing modality changed with the sample size. This insight has implications for planning large-scale neuroimaging studies, showing that results from small-scale pilot experiments can be misleading.

Our analyses comprised multiple brain imaging modalities (T1, rfMRI, DWI) and a wide range of sociodemographic, cognitive, behavioral, and mental health target phenotypes, evaluated using both linear and nonlinear machine learning frameworks on one of the largest available brain-imaging datasets. Due to the diversity of imaging modalities and target phenotypes, and due to sample sizes orders of magnitude beyond the size of traditional neuroimaging studies, we cautiously expect our results to generalize to other neuroimaging-based phenotype prediction scenarios.

Foundational for the present study is the premise that prediction performance follows strict mathematical laws. The gain in prediction accuracy that is enabled by an increase in sample size can be modeled and extrapolated. This allowed us to, in essence, forecast the prediction performance that we would likely reach at sample sizes orders of magnitude larger than the datasets of today. Learning curve extrapolation has already been applied in machine translation (Hestness et al., 2017), genomics (Hess & Wei, 2010; Mukherjee et al., 2003), and radiology (Cho et al., 2016) to assess sample size requirements. Regarding the estimation of learning curves, our investigation makes at least three key contributions. First, we demonstrated consistent and highly accurate (average goodness-of-fit R^2^=0.99) power law scaling of prediction accuracy with sample size in 48 (16 target phenotypes x 3 modalities) common neuroimaging data analysis scenarios. The diversity of analyzed phenotypes and imaging modalities suggests that the power law functional form can describe learning curves for neuroimaging data, independent of target phenotype or imaging modality. Second, we demonstrated that the underlying power law allows for extrapolating learning curves. While earlier studies (Cho et al., 2016; Hess & Wei, 2010) assumed the integrity of out-of-sample extrapolation without empirical evidence, we experimentally validated our ability to extrapolate learning curves based on subsamples of data. Finally, we conceptually linked the ceiling accuracy of extrapolated learning curves to the amount of predictive information contained in the data. Particularly for research in precision medicine, it is crucial to estimate whether neuroimaging data could ever be predictive of a certain disease or treatment response at a clinically useful accuracy. We argue that the ceiling accuracy of extrapolated learning curves can serve as an estimate of the maximally achievable accuracy, and partially answer the question of potential for clinical use. Our present study is, to the best of our knowledge, the first to use the theory of learning curves as an empirical tool to assess the irreducible error, or maximal accuracy, of real-world prediction tasks.

Will structural and functional neuroimaging benefit from exceedingly large sample sizes? In machine learning benchmarks datasets of comparable dimensionality, like MNIST (Lecun et al., 1998) or Fashion (Xiao et al., 2017), linear models approach saturation accuracy at around 1000 samples (Schulz et al., 2020). One may reasonably expect our neuroimaging data to follow a similar pattern. Indeed, for the prediction of sex and age in all modalities and the prediction of aspects of cognitive function T1 and DWI, linear models begin to saturate (Fig. 3). However, for nearly all other combinations of imaging modality and target phenotype, linear models appear to operate far below their ceiling accuracy. In conjunction with the observation that linear models are competitive with more complex nonlinear models at present sample sizes (Christodoulou et al., 2019; Dadi et al., 2019; Schulz et al., 2020; Dufumier et al., 2021), this allows for several conclusions.

Most importantly, there may be more predictive information contained in neuroimaging data than early small sample trials indicate. The extent to which quantifiable measures of behavior, cognition, even mental health can be inferred from structural or functional brain MRI is contentious in the quantitative neuroscience community (Hardcastle & Stewart, 2002; Logothetis, 2008; Uttal, 2011; Farah, 2014; Shifferman, 2015; Jonas & Kording, 2017; Falkai et al., 2018). For instance, in the ABCD challenge benchmarking the prediction of fluid intelligence based on T1 data, most contributing researchers reported low (R^2^<0.04) explained variance (Pohl et al., 2019). While some contributing researchers hypothesized insufficient sample size (Zhang-James et al., 2019), others speculated that “structural features alone do not contain enough information related to fluid intelligence to be useful in prediction contexts” (Guerdan et al., 2019). Our learning curve extrapolation suggests that the amount of explained variance could likely be doubled - or using rfMRI data even quadrupled - given enough training samples, and that we by far have not exhausted the predictive information in neuroimaging data. Though we cannot draw definitive conclusions on whether structural and functional MRI are operating on appropriate temporal or spatial resolutions, our results give cause for optimism that current spatiotemporal resolutions are “good enough”. The majority of medium-sample-regime HCP-sized (∼1k samples) studies on neuroimaging-based phenotype prediction have likely severely underestimated the accuracy which can be achieved in the limit of very large available sample sizes for predictive modeling.

Contrary to our expectations, rfMRI provided the best prediction accuracy in the limit of large sample sizes of many prediction tasks. Functional MRI is often criticized for being highly susceptible to a variety of noise sources (Liu, 2016; Raz et al., 2005), having low temporal resolution (Kim et al., 1997; Glover, 2011), and relying on the BOLD signal as a proxy for neural activity that is far removed from the actual local field potentials (Logothetis, 2008; Uttal, 2011). Such skepticism is, superficially, supported by comparably low prediction performance at small sample sizes (Fig. 2). However, learning curve extrapolation projected that rfMRI would eventually outperform structural imaging modalities in the prediction of the majority (9/16) of target phenotypes. Even with all its shortcomings, rfMRI appears to have a wealth of extractable predictive information. Given that fMRI likely leaves much room for technological innovation (Duyn, 2012; Patz et al., 2019), we conclude that fMRI could well allow for single subject prediction on very large datasets of the future.

Further, we observed that prediction performance for sex and age is saturating before all other phenotypes. In neuroimaging-based phenotype prediction, there is a well-founded fear that the model may mostly rely on confounding variables like sex and age to derive its prediction. Our results allow for careful optimism in this regard. Most phenotypes showed continuous improvements in achievable accuracy for the sample size ranges in which accuracies for age and sex confounds are already saturating. Thus, accuracy gains are unlikely to be driven primarily by age and sex confounds, and by exclusion principle more likely to be driven by phenotype-specific effects.

Finally, our results suggest that neuroimaging researchers may need to recalibrate what they consider large sample sizes. Average machine learning studies in the field of precision psychiatry include hundreds up to a few thousand samples (Arbabshirani et al., 2017); ten thousand is conventionally considered very large. However, one hundred thousand, even one million, samples may not be sufficient to characterize simple linear modes for most target phenotypes (Fig. 3). This is consistent with sample sizes of widely acknowledged reference datasets from computer vision, for example the popular Imagenet dataset, which comprises 14 million images. As brain images have an even higher dimensionality than photos from Imagenet (cubed instead of squared resolution), excessively large sample sizes may prove necessary to fully extract predictive information in neuroimaging data.

Our collection of findings underlines that “snapshot” measurements of accuracy at a single sample size can be highly misleading when deciding which imaging modality is most promising for predicting a given target phenotype. Particularly cognitive target phenotypes featured cross-over effects, where the most informative imaging modality changed with increasing sample size. Take the example of fluid intelligence (Fig. 2): Accuracy at a few hundred samples would suggest that T1 yields best performance, contains most information about the target, and should be prioritized for further research. At a few thousand samples, a researcher may conclude that all modalities work comparably well and that it barely matters which modality to prioritize for further data collection. Only when considering the extrapolated learning curve, did it become clear in the present investigation that rfMRI can be expected to substantially outperform structural modalities at high sample sizes for the prediction of fluid intelligence. This cross-over effect is likely driven by different phenotype-specific signal-to-noise ratios of different imaging modalities, leading to heterogeneous sample efficiency and, consequently, differently shaped learning curves.

Modality cross-over effects have implications for the planning and design of large-scale neuroimaging studies. Large-scale studies are often preceded by a pilot experiment to assess sample size requirements and to determine which particular neuroimaging techniques or modalities are most promising for the research question at hand. Without characterizing learning curves, pilot experiments will often produce deceptive results regarding the optimal imaging modality, like erroneously discarding rfMRI in favor of T1 in our example of fluid intelligence, leading to suboptimal design choices down the road. In contrast, learning curve extrapolation from the pilot experiment can reveal not only the optimal imaging modality in the limit of infinite samples but also for specific sample size regimes, facilitating superior design choices regarding sample size and imaging protocol of large-scale neuroimaging studies.

In the present study, mental health phenotypes could be predicted to a lesser accuracy compared to sociodemographic and cognitive phenotypes, alcohol and smoking behavior. This could be either due to particularly high noise in the features which are predictive for mental health compared to other phenotypes or due to comparatively high noise in the target variable. In our results, prediction appeared to work best for objective measures like age, reaction time, and fluid intelligence, and worst for noisy measures derived from subjective experience, like neuroticism or friendship satisfaction. Thus, we hypothesize that noise in the target variable may play an important role in the comparatively low prediction accuracy for mental health phenotypes.

This interpretation of our results ties into current discourse in psychiatry. Many diagnostic categories may insufficiently map on the underlying neurobiology (Hyman, 2007) or suffer from low inter-rater and low test-retest reliability (Kraemer et al., 2012; Regier et al., 2013). The concept of unsuitable labels provides a potential way to increase accuracy besides the collection of more and more data: Target phenotypes like depression or neuroticism (as used in this study) may be too noisy due to the subjectivity of individual experience or may insufficiently map on the underlying neurobiology and may need to be redesigned or split into constituent parts to optimize prediction.

The introduction of Research Domain Criteria (Insel et al., 2010) targets this problem by searching for “new ways of classifying mental disorders based on dimensions of observable behavior and neurobiological measures” (Cuthbert, 2015). Such newly derived phenotypes are expected to yield improved reliability and validity. Thus, Research Domain Criteria may not only improve clinical practice but also benefit neuroimaging-based precision psychiatry by improving the sample efficiency of machine learning models.

Our study has multiple conceptual and technical limitations. Conceptually, our learning curves inherently only give a lower bound to practically achievable prediction accuracy for a given model and a given representation of input data. A different model or a different representation of the input data may give different, potentially even better, results. While we cannot fully exclude such possibilities, we are confident in our results on both accounts. For the sample size range analyzed in this study (32 thousand and extrapolated up to one million), our learning curves may be quite close to the ground-truth achievable accuracy, even for arbitrarily expressive models (cf. Fig. 5). We give one empirical and one principled argument for this conjecture. Empirically, a number of recent studies found that linear models performed comparably to their more sophisticated nonlinear counterparts. In prior work (Schulz et al., 2020), we saw virtually no difference in performance when moving from linear models to kernel Support Vector Machines, Random Forests, Gradient Boosting, and Deep Neural Networks. Dufumier et al. (2021) confirmed this result and concluded that “simple linear models are on par with SOTA [state-of-the-art] CNN [convolutional neural networks] on VBM [voxel-based-morphometry], which suggests that DL [deep learning] models fail to capture non-linearities in the data”. Though these results are critically discussed by Abrol et al. (2021), it does remain controversial how much complex nonlinear models may improve over a well-tuned linear baseline in the analysis of neuroimaging data. High levels of noise in neuroimaging data may effectively linearize decision boundaries, potentially leaving little nonlinear structure for machine learning models to exploit (Schulz et al., 2020; Nozari et al., 2021). Even if the task of mapping a brain image to a phenotype is nonlinear, we have a principled reason to assume that this nonlinear predictive structure can rarely be exploited at present sample sizes. Our results show that linear models are still operating far below their ceiling accuracy at present sample sizes for nearly all of the analyzed target phenotypes. It follows that the parameters that constitute the linear model cannot be adequately estimated at present sample sizes. The linear model is the simplest possible mapping from features to prediction target, and any nonlinear extension requires additional parameters that would need to be estimated from the same insufficient data. We argue that if there is insufficient data to characterize a linear interaction, then there is little reason to expect to be able to characterize a more complex nonlinear interaction from the same data - unless the model implements an inductive bias specifically suited to the precise type of nonlinear interaction in the data, e.g., by incorporating neuroscientific domain knowledge.

Regarding a better representation of features, we are using the state-of-the-art representation of neuroimaging data, derived from years of experience and incorporating vast neuroscientific domain knowledge from nonlinear registration to feature creation based on cortical or volumetric parcellation. For a machine learning model like, for example, a deep neural network operating on minimally preprocessed T1 images to learn an internal representation that outcompetes the carefully hand-crafted representations we have available today is a substantive challenge, and positive results (Abrol et al., 2021; Peng et al., 2021; Plis et al., 2014; Vieira et al., 2017) are still controversially debated (He et al., 2020; Schulz et al., 2020; Dufumier et al., 2021).

Technical limitations pertain to the representation of target variables and to uncertainty quantification. It is difficult to make direct comparisons of prediction accuracy between different target phenotypes due to heterogeneous coding of the target variables. The explained variance (R^2^), which we report as a metric of prediction performance, has no intrinsic meaning and can only be interpreted in reference to the given coding, retest-reliability, and construct validity of a given target variable.

Finally, the quantification of uncertainties on the results of cross-validation schemes is an area of active research and has no established solutions. Our Monte Carlo cross-validation approach should yield legitimate estimates of uncertainty for small sample sizes, for which subsampled sets will be approximately statistically independent. When we approach 32 thousand training samples, sampled from a finite base population, statistical independence is violated, and error bars in all figures should be taken with caution. Consequently, we intentionally restrict ourselves to mostly qualitative interpretation of results and refrain from explicit statistical inference on the level of learning curves, and from reporting explicit confidence intervals on the results of learning curve extrapolations.

Taken together, we argue that the amount of predictive information contained in neuroimaging data, particularly in rfMRI, is likely underestimated, combined with, and partly due to, an overoptimistic assessment of the sample efficiency of machine learning models on neuroimaging data. We recommend characterizing and extrapolating learning curves as an essential part of pilot experiments to test feasibility by estimating the achievable accuracy, assess sample size requirements, and establish the optimal imaging modality regardless of cross-over effects.

## Methods

### Dataset and feature spaces

Our analyses required large sample sizes to reliably estimate learning curves. Hence, we based our analyses on the UK Biobank (Sudlow et al., 2015), which has been described as the “world’s largest multi-modal imaging study” (Littlejohns et al., 2020). The UK Biobank provides genotyping as well as extensive phenotyping data on approximately half a million participants, out of which 46197 (June 2021 release) underwent additional medical imaging. Structural T1-weighted brain images, resting-state fMRI, and diffusion-weighted brain images are available for each of these participants. For details on the UK Biobank’s data acquisition and processing protocols, please refer to Alfaro-Almagro et al. (2018).

A majority of past and present studies on machine learning for clinical neuroimaging rely on features designed by domain experts; cf. recent reviews on machine learning in epilepsy (Sone & Beheshti, 2021), autism (Xu et al., 2021), stroke (Sirsat et al., 2020), and mild cognitive impairment (Ansart et al., 2021). The UK Biobank directly provides widely used feature representations (IDPs) for structural, functional, and diffusion tensor imaging. To align with common usage of neuroimaging feature representations and to increase reproducibility, we used the UK Biobank-provided features for our analysis. The structural MRI features (1425 descriptors) represented regional gray and white matter volumes, and parcellated cortical thickness and surface area (Alfaro-Almagro et al., 2018). Resting-state functional MRI was distilled into a functional connectivity matrix (1485 descriptors), based on networks derived from a 100-component group ICA (Alfaro-Almagro et al., 2018). Diffusion-weighted imaging features (675 descriptors) represented fractional anisotropy, mean diffusivity and tensor mode, intra-cellular volume fraction, isotropic or free water volume fraction and orientation dispersion index for “over 75 different white-matter tract regions based both on subject-specific tractography and from population-average white matter masks” (Miller et al., 2016). For details on the UK Biobank’s pre-computed feature representations, please refer to Alfaro-Almagro et al. (2018). Aside from standard scaling, features were used exactly as provided by the UK Biobank.

### Prediction targets and target variable coding

Widely studied sociodemographic, cognitive, behavioral, and mental health phenotypes served as prediction targets for our analyses. Specifically, we included participant age (UK Biobank field-ID 21003-2), sex (field 31-0), education score (field 26414-0), and household size (number of people in household, field 709-2) as sociodemographic phenotypes. Fluid intelligence (summary score, field 20016-2), reaction time (mean time to correctly identify matches, field 20023-2), numeric memory (maximum digits remembered correctly, field 4282-2), trail making (interval between previous point and current one in alphanumeric path, field 6773-2) represented cognitive phenotypes. Alcohol consumption (intake frequency, field 1558-2), tobacco consumption (intake frequency, field 1249-2), physical activity (International Physical Activity Questionnaire activity group, 22032-0), TV consumption (hours per day, field 1070-2) represented behavioral and lifestyle phenotypes. Finally, depression (ever felt depressed for a whole week, field 4598-2), neuroticism (summary score, field 20127-2), friendship satisfaction (field 4570-2), and financial satisfaction (field 4581-2) represent mental health. For detailed information on the prediction targets, please refer to the UK Biobank online documentation (https://biobank.ndph.ox.ac.uk/showcase).

With the exception of sex and depression, all targets are either continuous or ordinally represented and were treated in a regression setting. Prediction of sex and depression, both binary targets, was treated as a classification task. All target phenotypes were provided by UK Biobank and used as-is, excluding “prefer not to answer” and “do not know” responses on a per-phenotype basis. Full phenotype information was not available for all participants, so that we generally report results for 16 thousand participants or the maximum available sample size per target phenotype.

### Machine learning models and out-of-sample validation

For each combination of MRI modality, target phenotype, and training sample size, we subsampled the data into a training set (n=256, 362, 512, …, 32k), a validation set for hyperparameter tuning (n=2k), and a test set for final evaluation of prediction accuracy (n=2k). The subsampling into train, validation, and test sets was repeated 20 times (Monte Carlo cross-validation, also known as repeated random sub-sampling validation) to provide an averaged accuracy as well as uncertainty estimates. Uncertainties were derived by bootstrapping over the cross-validation resamplings. Samples of train, validation, and test sets can be considered approximately independent for small train set sizes. For large train set sizes, train sets will overlap, and uncertainty estimates must be viewed with caution.

A new machine learning model was trained on the train set, tuned on the validation set, and the best hyperparameter configuration evaluated on the test set, for each MRI modality (n=3), target phenotype (n=16), training sample size (n=15), and cross-validation resampling (n=20). We used ridge regression (l2 regularized linear regression, sklearn.linear_model.Ridge, Pedregosa et al., 2011) for regression tasks, and logistic regression (l2 regularized, sklearn.linear_model.LogisticRegression, Pedregosa et al., 2011) for classification tasks. Regularized linear models are considered highly competitive in the analysis of neuroimaging data and performed on par with more complex nonlinear models, such as kernel support vector machines and deep artificial neural networks, on similar prediction tasks (Dadi et al., 2019; Schulz et al., 2020; Dufumier et al., 2021).

Hyperparameter tuning on the validation set was performed separately for each fitted model. For ridge regression, alpha values from 2^-15 to 2^15 were evaluated in steps of 2^-(2n+1). For logistic regression, l2 penalty weights ranged from 2^-20 to 2^10 in insteps of 2^-(2n). Hyperparameter ranges and granularity of the search range were checked visually for each modality x target phenotype combination.

### Learning curve fitting

The empirical scaling of prediction performance with increasing training sample size is called a learning curve. Theoretical results from statistical learning theory state that learning curve bounds follow a power law function (Amari, 1993; Amari et al., 1992; Haussler et al., 1996; Hutter, 2021). Empirically, power law scaling was shown for models ranging from linear estimators to deep neural networks (Cortes et al., 1994; Hestness et al., 2017). Thus, we fitted the expected power law functional form [*αn*^−β^ + *γ*] to our empirical data. Learning curves were averaged over the 20 cross-validation resamplings before fitting the power law via nonlinear least squares (scipy.optimize.curve_fit, Virtanen et al., 2020). Parameters were bound to 0<α<inf, -inf<β<0, and 0<γ<1. Uncertainty estimates were derived via bootstrapping over the cross-validation resamplings.

## Code and data availability

For efficient calculation of learning curves, we used the Empirical Sample Complexity Estimator (https://github.com/maschulz/FIXME). Analysis and visualization code is available at https://github.com/maschulz/FIXME. Neuroimaging data was obtained from UK Biobank under Data Access Application 33073 and are available on request directly from the UK Biobank (http://www.ukbiobank.ac.uk/register-apply/).

## Acknowledgments

We thank Moritz Seiler, Matt Chapman-Rounds, and Braedon Lehman for insightful discussions and feedback on the manuscript. We thank the UK Biobank participants for their voluntary commitment and the UKBiobank team for their work in collecting, processing, and disseminating these data for analysis. This research was conducted using the UK Biobank Resource under project-ID 33073. Computation has been performed on the HPC for Research cluster of the Berlin Institute of Health. The project was funded by the Deutsche Forschungsgemeinschaft (DFG, German Research Foundation) under project-ID 414984028 - CRC 1404. D.B. was supported by the Healthy Brains Healthy Lives initiative (Canada First Research Excellence fund), the CIFAR Artificial Intelligence Chairs program (Canada Institute for Advanced Research), Google (Research Award), and by NIH grant R01AG068563A. S.H. acknowledges funding from the European Research Council (ERC) under the European Union’s Horizon 2020 research and innovation program (Grant agreement No. 758985). J.D.H. was funded by the DFG EXC 2002/1 “Science of Intelligence” – project-ID 390523135. K.R. was supported by the DFG (389563835; 402170461 - TRR 265; 414984028 - CRC 1404; 442075332 - RU 5187) the Brain & Behavior Research Foundation (NARSAD young investigator grant) and a DMSG research award.

